# Complex motion trajectories are represented by a population code from the ensemble activity of multiple motion-sensitive descending interneurons in locusts

**DOI:** 10.1101/2024.09.29.615702

**Authors:** Sinan Zhang, John R. Gray

**Author notes:** Corresponding author Correspondence: Sinan Zhang Department of Medical Genetics, Cumming School of Medicine, University of Calgary, Calgary, Alberta, Canada T2N4N1.

## Abstract

Adaptive locust flight relies on rapid detection and processing of objects moving in the visual field. One identified neural pathway, comprised of the lobula giant movement detector (LGMD) and the descending contralateral movement detector (DCMD), responds preferentially to approaching objects. The LGMD receives retinotopic inputs from ipsilateral ommatidia and generates spikes in a 1:1 ratio in the DCMD, which synapses contralaterally with multiple locomotion-related neurons. Other motion-sensitive neurons have also been identified in locusts but are not as well characterized. To better understand how locusts process visual information, we used multichannel neural recordings within a stimulus arena and presented various complex visual stimuli to investigate the neural encoding of complex object motion. We found multiple discriminated units that responded uniquely to visual motion and categorized the responses. More units responded to motion trajectories with a looming component, compared to translations across the visual field. Dynamic factor analysis (DFA) of discriminated units revealed common trends that reflect the activity of neural ensembles. The numbers and types of common trends varied among motion trajectories. These results increase the understanding of complex visual motion processing in this tractable system.

## Introduction

Animals orienting in natural environments need to detect salient sensory cues and rapidly respond with appropriate behaviours. This is important in many scenarios, such as locating food sources, finding mates, and avoiding approaching predators. In many species, such as monkeys (Cooke and Graziano 2004), cats (Regan et al. 1979); pigeons (Wang and Frost 1992; Wu et al. 2005), crabs (Oliva et al. 2007; Medan et al. 2015), and butterflies (Céchetto et al. 2022), motion-sensitive visual neurons have been identified, and shown to be critical in initiating avoidance behaviours.

Compared to vertebrates, invertebrates possess relatively tractable nervous systems, yet are still capable of performing complex adaptive behaviours in response to external stimuli. An example is the collision avoidance behaviour in the migratory locust, *Locusta migratoria*. Locusts migrate in large swarms, comprised of thousands of individuals (Uvarov 1977). Field observations found that they rarely collide with conspecifics and, in the meantime, can still avoid predators (Rind and Santer 2004). In locusts, a neural pathway, comprised of the lobula giant movement detector (LGMD) and the descending contralateral movement detector (DCMD), has been identified and thoroughly studied. The LGMD receives inputs from an entire compound eye on the ipsilateral side (O’Shea and Rowell 1976), and signals the DCMD to generate action potentials at a 1:1 ratio (Rind 1984). The DCMD axon, as indicated by the name, descends towards the contralateral nerve cord. The LGMD/DCMD pathway is tuned to preferentially detect objects approaching along a collision course by increasing its firing rate that peaks near the projected time of collision (TOC) (Gabbiani et al. 1999, 2001, 2002; Guest and Gray 2006; McMillan and Gray 2012; Dick and Gray 2014). Parameters of LGMD/DCMD responses, including the peak time, peak firing rate, and the firing rate when the object reaches a specific angular threshold, have been associated with the property of object motion (Gabbiani et al. 1999; Dick and Gray 2014), as well as the timing of collision avoidance behaviours (Santer et al. 2006). It is noteworthy that the DCMD peak time is likely not the cue to initiate behaviour, since the peak time can occur after TOC (Rind and Santer 2004).

However, although the response of the LGMD/DCMD pathway is relatively consistent, the behavioural response of flying locusts can be unpredictable (Dawson et al. 2004; Santer et al. 2006; McMillan et al. 2013; Chan and Gabbiani 2013). Flying locusts respond to the same looming stimuli with a wide range of behaviours, including steering, gliding, and stopping, suggesting that the DCMD response needs to be appropriately timed within the wing beat cycle to generate specific behaviours (Santer et al. 2006), and LGMD/DCMD is likely not the sole pathway that mediates motion-evoked behaviours. Indeed, other motion-sensitive neurons have been identified in locusts, such as the LDCMD (Gray et al. 2010) and the columnar and tangential neurons in the central complex (Rosner and Homberg 2013). Many of these neurons are sensitive to looming stimuli, yet they also show distinct response preferences, such as sensitivity to small objects or directional selectivity. In addition, although unable to form focused retinal images, the ocelli are responsive to large-field luminance changes, which can result from a large looming object, and therefore the ocellar neurons can also contribute to the encoding and generation of avoidance behaviours (Taylor 1981). Multichannel recordings revealed that a larger number of units (neurons), on average 10-30 units per locust, are responsive to object motion (Dick et al. 2017; Parkinson et al. 2020). These units can be categorized based on response types, some of which resemble the activity of previously identified motion-sensitive neurons. However, the encoding of complex motion by the neuronal population is not yet fully understood. Two studies that recorded from multiple locust visual interneurons neurons (Dick et al. 2017; Parkinson et al. 2020) presented a 7 cm black disc at a constant velocity of 300 cm s^-1^ (*l*/|*v*| = 11.7 ms, where *l* is the half size of the object and |*v*| is the absolute velocity). No previous studies had explored multiunit responses to stimuli travelling at different values of *l*/|*v*|.

Neuronal populations, commonly characterized as neural ensembles, are groups of neurons that are involved in a common neural computational task (Panzeri et al. 2015). Combined activity of multiple neurons can provide a more comprehensive interpretation of the perceived sensory inputs, and better guide behavioural responses. Using multichannel electrodes, synchronized activity of multiple neurons can be recorded and decoded in a mathematical approach, which can be related to the timing of behaviours. In dragonflies, a group of 16 target-selective descending neurons encode a population vector that describes the orientation of the target (Gonzalez-Bellido et al. 2013). During visual navigation of *Drosophila*, population coding has also been observed via calcium imaging (Seelig and Jayaraman 2015).

The central nervous system of a locust is comprised of a brain, which processes sensory information, and a segmented ventral nerve cord. Each segment of the ventral nerve cord is controlled by a ganglion. From anterior to posterior, the series of ganglia includes the suboesophageal, prothoracic, mesothoracic, metathoracic, seven abdominal ganglia, and the terminal ganglion (Burrows 1996). In the collision avoidance behaviour of locusts, the processed visual information is passed downstream to corresponding motor neurons that control the wings and legs, which are located in the mesothoracic and metathoracic ganglia (Burrows and Rowell 1973; Simmons 1980; Pearson et al. 1985; Boyan 1989). Therefore, the axons of motion-sensitive descending interneurons that are involved in processing visual motion, including the DCMD, go through the connectives between the brain and the mesothoracic ganglion. Here, using multichannel extracellular recordings anterior to the prothoracic ganglion, we simultaneously recorded from multiple neurons when presenting visual stimuli to rigidly-tethered locusts. After spike sorting and unit discrimination, units that showed significant change in firing rate during the presentation of visual stimuli were identified. We found that multiple units responded to visual motion, yet the response type varied. More units responded to looms and compound trajectories that contain a looming phase, compared to simple translations. Common trends, which represent neural ensemble activity, were extracted from the population activity, and revealed that most units responded to loom with a peak near TOC. The population response was also tuned by motion trajectory and velocity.

## Materials and Methods

### Animals

Nineteen gregarious adult male locusts (*Locusta migratoria*) were selected from a crowded colony maintained at the University of Saskatchewan, Canada and provided with a diet consisting of wheat grass and bran, and housed in a controlled environment with a 12-hour light and 12-hour dark cycle at 30 °C. We used locusts that were at least three weeks past the imaginal moult.

### Preparation

Before the experiments, the locust’s legs were removed, and a rigid tether was attached to the ventral surface of the thorax using melted beeswax. To expose the paired connectives of the ventral nerve, a portion of the ventral cervical cuticle was excised, anterior to the prothoracic ganglion. The prepared locust was then transferred to a stimulus arena (Fig 1.a, (Gray et al. 2002). All experiments were conducted at an ambient temperature of approximately 25 °C.

**Figure 1.**
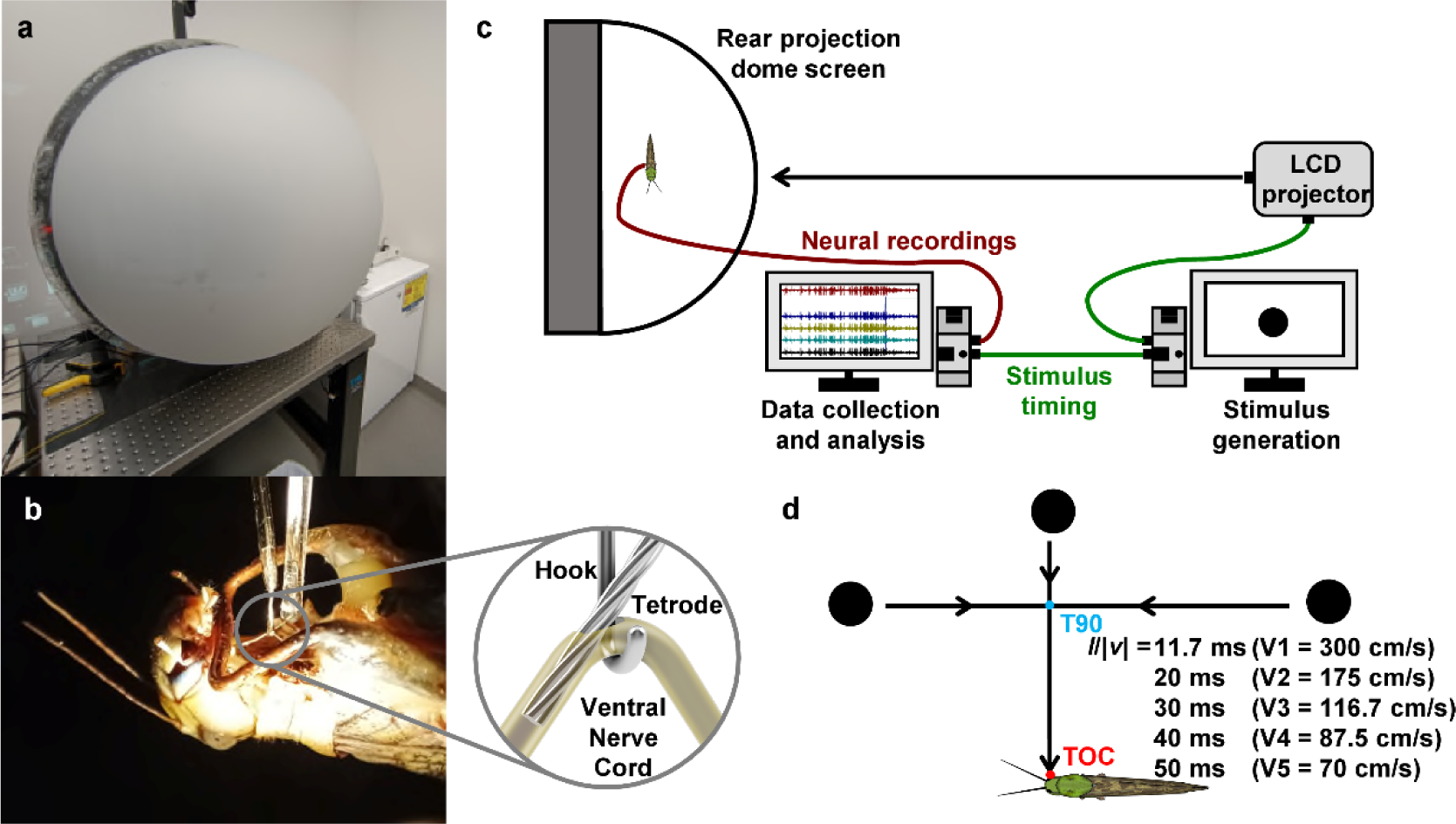
Experimental setup and visual stimuli. a) The flight simulator with the dome screen closed. b) The preparation with demonstration of the tetrode insertion method (inset). The ventral nerve cord was lifted with a silver hook electrode, and the tetrode was inserted parallel through an opening of the protective sheath. c) Diagram of the experimental setup. Visual stimuli were rendered and projected onto the dome screen while neural activity was recorded simultaneously with the multichannel electrode. A 5-V stimulus timing pulse was used to align the neural recordings with stimulus timing. d) Visual stimulus trajectories used in the experiment. All visual stimuli consisted of a 7 cm diameter black disc travelling against a white background. A total of five trajectories were used, including direct loom, translation (anterior towards posterior, or posterior towards anterior), and compound trajectories that started as a translating movement and transitioned into a direct loom at a 90°azimuthal angle. The time of the projected collision (TOC) and the time that the object passed through 90°azimuth (T90) were used to align recordings from different trajectories. For each trajectory, we used five velocities. The *l*/|*v*| values shown represent the half size of the object divided by the absolute value of the velocity. The order of all 25 visual stimuli was randomized for each animal.

We used a silver wire hook electrode to record whole nerve chord activity and stabilize the chord for tetrode insertion. Subsequently, we used a sharp glass electrode to create a small opening in the protective sheath surrounding the nerve cord. A twisted wire tetrode (Guo et al. 2014) was carefully inserted into the left nerve cord, anterior to the hook electrode (Fig 1.b). The tetrode consisted of four wires (diameter = 12.7 μm each, Model R0800, Sandvik-Kanthal Precision Technology, Hallstahammar, Sweden) that were twisted together and fused at one end, while the other end was separated and soldered to four channels of an adapter. The fused end of the tetrode was secured within a capillary tube for stability. A micromanipulator was used to maneuver the capillary tube, lowering the fused end of the tetrode into the nerve cord, while the other end (adapter) was connected to four channels of a differential amplifier (Model 1700, A-M Systems, United States). A silver ground wire was inserted into the locust’s abdomen.

After confirming a low signal-to-noise ratio of the recording by observing the neural response to a waving hand, petroleum jelly was applied around the electrodes and the nerve cords to insulate the recording site from the hemolymph and prevent desiccation. The entire experimental setup was then rotated 180°to position the locust in a dorsal-side up orientation at the center of a rear projection dome screen. The longitudinal axis of the locust was arranged perpendicular to the apex of the dome, with a distance of 24 cm between the locust’s head and the screen (Fig 1.c). In this configuration, we designated the front of the locust as 0°azimuth, the rear as 180°azimuth, and the perpendicular direction as 90°azimuth (Fig 1.d).

### Visual stimuli

Visual stimuli were generated at a resolution of 1024×768 using a Python-based program EVSG (Stott et al. 2018). The program’s code included correctional factors to compensate for the curvature of the dome. Each pixel on the screen was ∼0.7×0.7 mm^2^, corresponding to a visual subtense angle of ∼0.4°. This subtense angle of individual pixels was below the angular resolution of locusts’ eyes (1°) (Horridge 1978). Individual stimulus frames were rendered in real-time by a GeForce GTX660 video card (NVIDIA Corporation, Santa Clara, United States) at a frame rate > 150 frames·s^-1^ and projected onto the dome screen using an InFocus DepthQ projector (InFocus, Portland, United States).

For each trial, we presented one of five different trajectories of visual stimuli, each travelling at one of five different velocities (Fig 1.d). The trajectories were derived from a subset of trajectories used in (McMillan and Gray 2012). All visual stimuli consisted of a 7-cm diameter black disc moving against a white background (Michelson contrast ratio = 0.98). The first trajectory was a simple loom approaching from 90°azimuth (Loom). The second and third trajectories were translations parallel to the longitudinal axis of the locust body, positioned at a virtual distance of 80 cm from the eye. Translations moved either from posterior to anterior (trans P-A) or from anterior to the posterior (trans A-P). The last two trajectories began as translations (in either direction) and transitioned to looming when at 90°azimuth (comp P-A and comp A-P). Each trajectory was presented at velocities of 300, 175, 116.7, 87.5, and 70 cm·s^-1^ (V1-5, respectively). The *l*/|*v*| value, which represents the ratio of the object’s half size (*l*) to the absolute velocity (|*v*|), is commonly used to describe looming objects and correlates to DCMD firing properties (Gabbiani et al. 1999; Dick and Gray 2014). For the 7-cm diameter disc, the five velocities corresponded to *l*/|*v*| values of 11.7, 20, 30, 40, and 50 ms, respectively. This range of *l*/|*v*| values has been previously used (Gabbiani et al. 2002; Dick and Gray 2014) and is biologically relevant (Santer et al. 2012). All visual stimuli were presented at an elevation of 0°.

The presentation sequence of all 25 visual stimuli was uniquely randomized for each locust. To prevent neural habituation, a 3-minute interval was maintained between presentations. Additionally, a direct loom was presented before and after the sequence to confirm that no attenuation occurred during the length of the experiment. A 5V pulse was generated at either the time that the object passed through 90°azimuth (T90) and/or the projected time of collision (TOC), both of which were automatically calculated based on the motion trajectory. This pulse was used to align neurophysiological data with the stimulus.

### Data Acquisition

Neural signals from all five channels, including one channel from the silver hook electrode and four channels from the twisted wire tetrode, were amplified using high-pass and low-pass filters set at 300 Hz and 5 kHz, respectively, and had a gain of 100×. The amplified neural signals, along with the 5V stimulus pulse, were then digitized using a USB data acquisition board (DT9818-OEM, TechmaTron Instrument Inc., Laval, Canada) and recorded at a sampling rate of 25 kHz/channel using DataView version 11 (W.J. Heitler, University of St Andrews, St Andrews, Scotland).

### Spike Sorting

Raw recordings were initially merged chronologically into a single file. Subsequently, DataView was used to apply a Finite Impulse Response (FIR) filter with a bandpass range of 500-2000 Hz and a Blackman window type. The filtered data from the four channels of the tetrode were then exported to Offline Sorter version 4.6 (Plexon Inc., Dallas, United States) for waveform detection and spike sorting.

The threshold for waveform detection was set at three times the standard deviation from the mean for each channel. Spike sorting was performed using a semi-automatic method based on the T-Dist E-M algorithm (Shoham et al. 2003). The degree-of-freedom (DOF) multiplier was set to 3, and the number of initial units was set to 26. Initial sorting in the three-dimensional (3D) feature space was carried out automatically, followed by manual discrimination of units based on waveform shape and amplitude (Fig 2.a-b). Multivariate Analysis of Variance (MANOVA) was used to assess whether the sorted units were statistically distinct from each other in the 3D cluster space (Supplementary Table 1).

**Figure 2.**
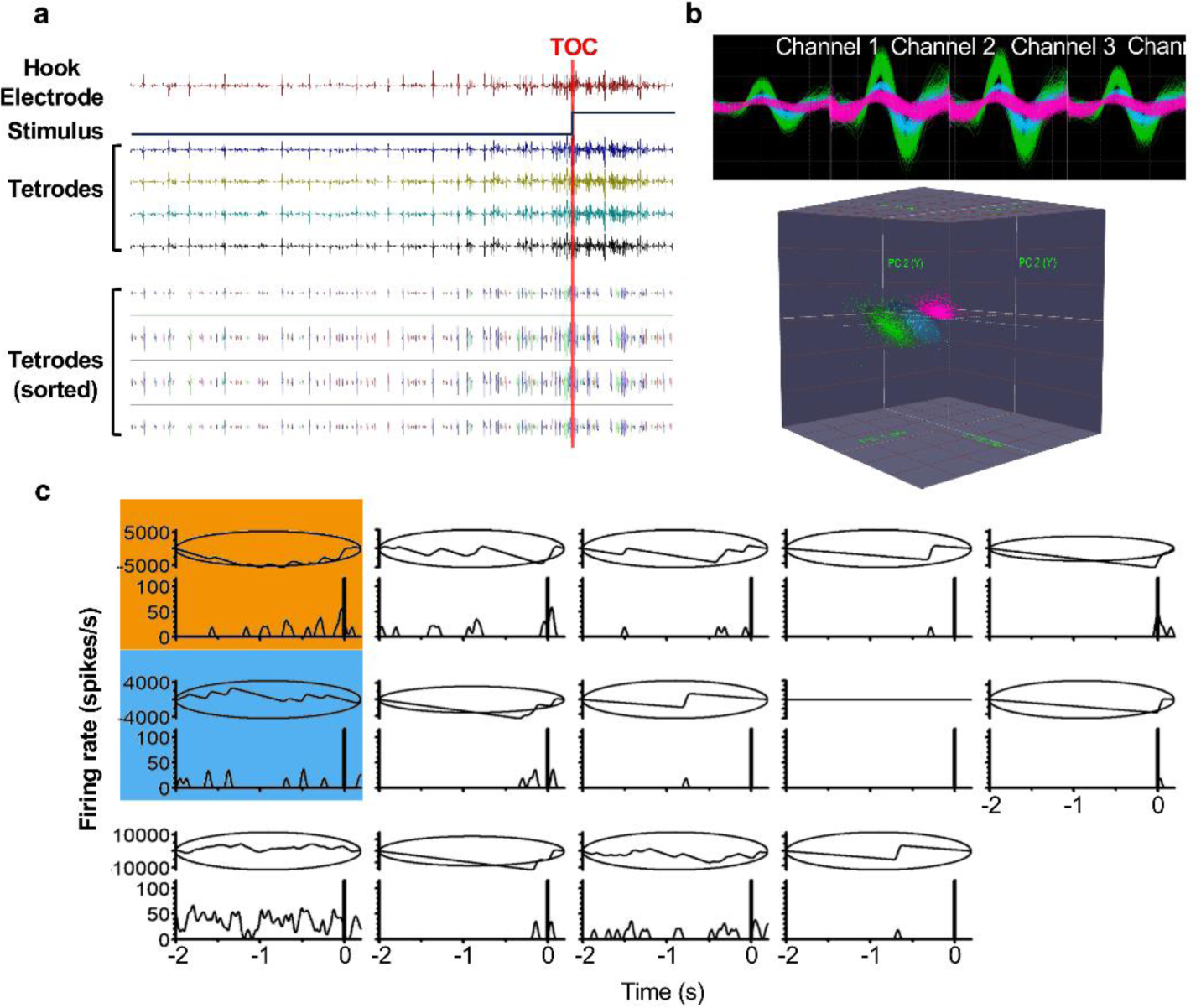
Spike sorting and identification of responsive units. a) Raw recording from the hook electrode, the stimulus event pulse, and the tetrode are shown on the top. Waveforms were detected using average plus three times the standard deviation as the threshold and sorted based on the cluster variance in the 3-dimentional (3D) feature space. Spike sorting results from the same recording time window (tetrodes sorted) is shown underneath the raw recordings. Each colour represents a different unit. b) Overlapped waveforms of three units recorded from all four channels of the tetrode (top), and the distribution of the same units in the 3D feature space (bottom). c) Identification of responsive units. For each discriminated unit, the peristimulus time histogram (PSTH), representing the change of firing rate during the stimulus presentation, is shown on the bottom of each block. The cumulative sum and 99% confidence level (represented by the ellipses) are plotted on the top half of each block. If the cumulative sum (represented by the line inside the ellipse) extended outside the ellipse (example with orange background), we designated the unit as responding. If the cumulative sum remained within the ellipse for the entire duration (example with blue background), we designated the unit as not responding.

### Spike Train Analysis

Spike times of the discriminated units were exported to NeuroExplorer version 5 (Plexon Inc., Dallas, United States). Peristimulus time histograms (PSTHs) were constructed using a 1-ms bin width and smoothed with a 50-ms Gaussian filter (Guest and Gray 2006). All trials were aligned either to the projected time of collision (TOC) or the time that the object passed through 90°azimuth (T90). For analysis, we designated trials to start when the object reached a subtense angle of 1°, which was the estimated time when the stimulus could be first detected by the locust, and ended 0.2 seconds after the TOC (Yakubowski et al. 2018; Stott et al. 2018). For translating trajectories, the trials began when the subtense angle exceeded 1°and ended when the subtense angle decreased below 1°.

The responsiveness of individual units was assessed by plotting the cumulative sum and a 99% confidence level ellipse (Fig 2.c). Units for which the cumulative sums touched or expanded beyond the ellipse were designated as responding to the visual stimulus (Ritzmann et al. 2008; Dick et al. 2017; Parkinson et al. 2020). Units for which the cumulative sum was contained within the ellipse were designated as not responding and were excluded from further analysis.

### Dimensionality reduction

The response of individual units (designated by different letters) was first examined to determine how many types of visual stimuli each unit responded to. Since the positioning of the tetrode may have differed slightly in each animal, we could not confirm that we recorded from the exactly the same neurons in each locust. Therefore, responsive units were pooled across all animals based on the visual stimulus and categorized into response types based on properties of the peristimulus time histograms (PSTHs). This provided a general description of the different types of observed responses. The categories were adopted from previous studies (Dick et al. 2017; Parkinson et al. 2020).

Dynamic Factor Analysis (DFA) was then used to extract common trends from the responsive units for each visual stimulus (Zuur et al. 2003; Dick et al. 2017). In this model, a set of observed time series (**y**) (units) is explained by a set of hidden random walks (**x**) (common trends) through linear combinations, represented by factor loadings (**Z**) and offsets (**a**). The model equations can be written as follows:

**y**_t_ = **Z** ×**x**_t_ + **a** + **v**_t_, where **v**_t_ ∼ MVN (0, **R**)

**x**_t_ = **x**_(t-1)_ + **w**_t_, where **w**_t_ ∼ MVN (0, **Q**)

In the model, the error terms of hidden processes (**x**) (common trends) and observations (**y**) (units) are represented by **w**_t_ and **v**_t_, respectively. Both **w**_t_ and **v**_t_ follow a multivariate normal distribution (MVN), with a mean of 0 and covariate metrices of **R** and **Q**, respectively. To identify the model, some constraints were applied: the first *m* (the number of hidden processes) elements of **a** were set to 0; **z**_i,j_ was set to 0 if j > i for the first *m*-1 rows of **Z**; **Q** was set to the identity matrix **I**_m_.

DFA was performed using the MARSS package version 3.11.4 (Holmes et al. 2012) in R 4.1.3 (R Core Team, 2022), using the Broyden-Fletcher-Goldfarb-Shanno (BFGS) method. We evaluated the model quality using the Akaike Information Criterion with a correction for small sample size (AICc) (Burnham and Anderson 2002). For each visual stimulus, DFA was iteratively performed, starting with one common trend. The number of common trends gradually increased until the AICc began to increase, and the model with the lowest AICc was selected as the optimal model.

Factor loadings (**Z**) of each common trend were used to normalize the range of common trends and rectify any potentially reversed trends. According to the DFA model, **y**_t_ = **Z** ×**x**_t_ + **a** + **v**_t_, for the *i-th* common trend ((**x**^T^)_i_), if the factor loadings of this common trend (**Z**_i_) were mostly negative, reversing the sign of (**x**^T^)_i_ and **Z**_i_ simultaneously would not impact the resulting observations (**y**). Additionally, this will make the direction of the common trend consistent with resulting observations, and thus provide a more accurate representation of the majority of unit activities. Similarly, the factor loadings of each common trend were scaled to (-1 to +1). We calculated the largest absolute value of the factor loadings for each common trend, divided all factor loadings by this value, and, correspondingly, multiplied the common trend with the same value. As discussed above, this will not affect the model fit, and can provide a comparable scale among different common trends. For common trends showing clear peaks, various parameters, such as peak time (when the firing rate reached peak firing rate), half-peak time (when the firing rate reaches 50% of the peak firing rate), subtense angle (SA) of the stimulus at peak and half-peak, peak width at half height (PWHH), half-peak-to-peak time, and peak-to-half-peak time, were measured from the PSTHs using R 4.1.3. These parameters were then compared between different stimuli.

### Statistical analysis

Statistical analyses were conducted using SigmaPlot version 12.5 (Systat Software, San Jose, USA) or R 4.1.3 (R Core Team, 2022), and the resulting plots were created with SigmaPlot. Prior to analysis, data were assessed for normality and equal variance using the Shapiro-Wilk test. Parametric data were described using the mean and standard deviation and presented using column graphs. Nonparametric data were described using the median and quartiles and presented using box plots.

For comparisons between unrelated variables, either one-way ANOVA (parametric) or Kruskal-Wallis One Way Analysis of Variance on Ranks (nonparametric) were utilized. Comparisons between related variables were performed using either one-way repeated measures (RM) ANOVA (parametric) or Friedman Repeated Measures Analysis of Variance on Ranks (nonparametric). Two-way comparisons of related variables were conducted using two-way RM ANOVA, followed by the Holm-Sidak post-hoc test when necessary. Linear correlations were fitted between each PSTH parameter and the *l*/|*v*| value. All statistical tests were two-tailed, and the significance level (*α*) was set at 0.05.

## Results

### Unit discrimination

For each animal, all 25 presentations (trials) were merged in chronological order for spike discrimination and unit sorting. Using a positive threshold set at three times the standard deviation over the mean, a total of 789,028 spikes were discriminated across all 19 animals, with an average of 41,527.8 spikes per locust and a standard deviation of 10,820.7 spikes per locust. These spikes were further sorted into 352 units in total, with an average of 22.9 units per locust and a standard deviation of 1.7 units per locust. MANOVA analysis confirmed that these units were statistically distinct in the 3D cluster space (*p* < 0.001) (Fig 2.b). Supplementary Table 1 provides a summary of the discriminated spikes and units from all 19 animals.

### Unit response

The responsiveness of each unit was individually examined with respect to all 25 unique visual stimuli. Supplementary Table 2 displays the responses of all discriminated units from one animal, with a total of 22 designated units (a-v). Each unit responded to at least one of the visual stimuli, ranging from a minimum of 2 responses (unit v) to a maximum of 23 responses (unit k). Twenty units responded to at least one of the simple looming stimuli at V1-V5. Units n and v did not respond to any looming stimuli at any velocity tested. All units responded to at least one of the ten translating trajectories (trans P-A or trans A-P at V1-V5) and one of the ten compound trajectories (comp P-A or comp A-P at V1-V5). Fig 3.a provides a summary of the number of stimuli each unit responded to within each trajectory category from the same animal. Out of the five velocities presented along each trajectory, individual units from this animal responded to a larger proportion of looming and compound trajectories, compared to translating trajectories (One-way repeated measure (RM) ANOVA, F_4_ = 9.356, *p* < 0.001), except for comp P-A vs. trans A-P (Holm-Sidak post-hoc test, *p* < 0.05) (Fig 3.b). Although there was no statistical difference, individual units, on average, responded to a larger proportion of A-P trajectories, compared to P-A trajectories, which aligns with previous findings that the LGMD/DCMD pathway is more sensitive to movements originating from the anterior visual field, compared to the posterior visual field (Rogers et al. 2010; McMillan and Gray 2012).

**Figure 3.**
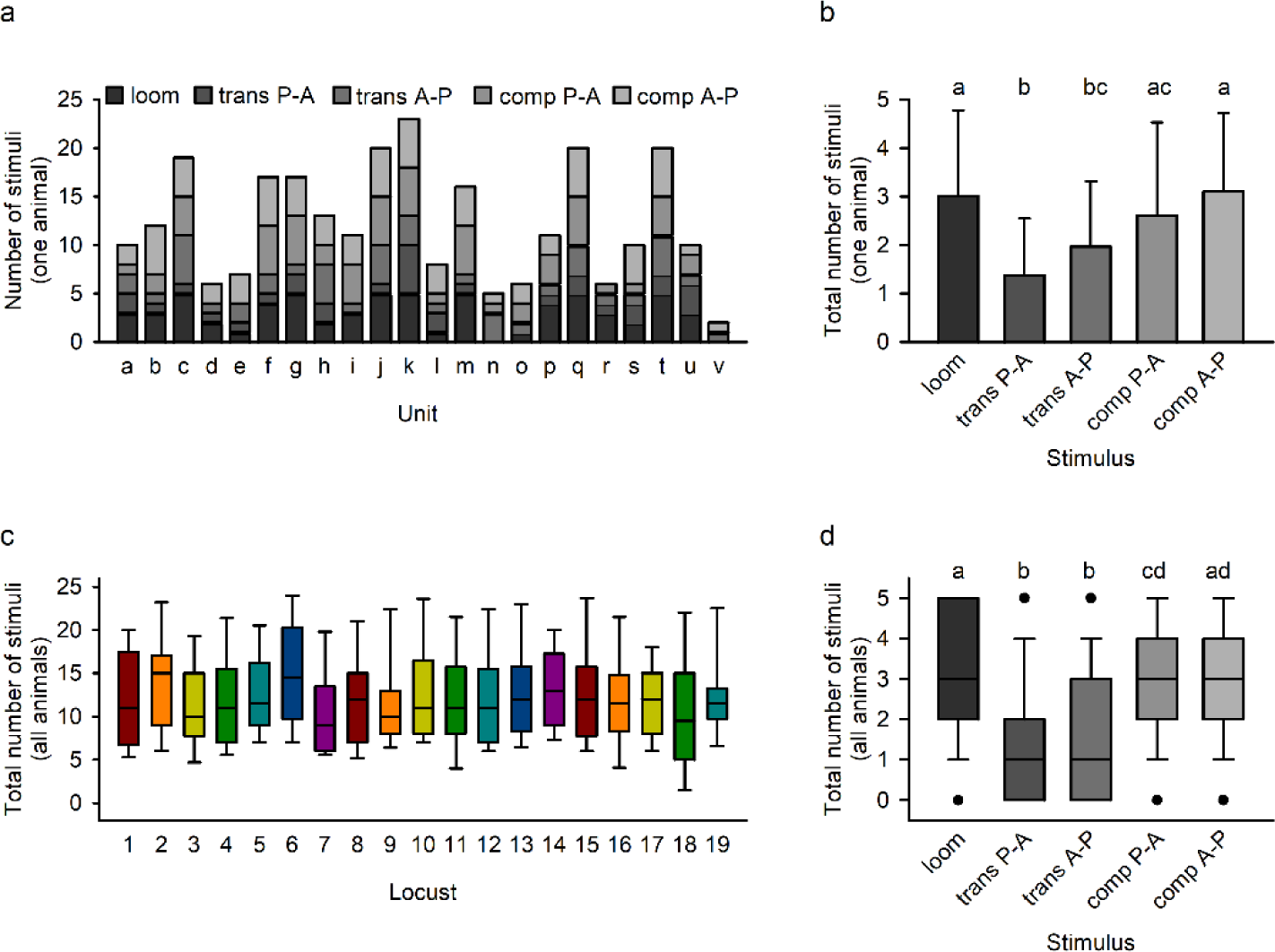
Comparison of the number of responding units for each stimulus trajectory. a) The response of all discriminated units from one animal. Each column represents the number of stimuli a unit responded to, and the shades represent the stimulus category. Each unit responded to at least one of the translating stimuli (trans P-A or trans A-P) and one of the compound trajectories (comp P-A or comp A-P). b) Column graph (mean + standard deviation) comparing the unit responses from the same animal in a. Letters abovethe columns represent statistical differences. c) The total number of stimuli individual units responded to across all 19 animals. The horizontal line within each box represents the median, while the lower and upper boundaries, the whiskers, and the dots represent the 25/75%, 10/90%, and 5/95% percentiles, respectively. d) Unit responses from all 19 animals. Within each box, the horizontal line represents the median, while the lower and upper boundaries, the whiskers, and the dots represent the 25/75%, 10/90%, and 5/95% percentiles, respectively. Letters above the columns represent statistical differences.

Considering all 436 units discriminated across the 19 animals, there was no significant difference in the number of stimuli each individual unit responded to (Kruskal-Wallis One Way Analysis of Variance on Ranks, H_18_ = 14.561, *p* = 0.692) (Fig 3.c), indicating unit consistency across recordings from different animals. When comparing responses to different trajectories, out of the five velocities along each trajectory, designated units from all locusts responded to a larger proportion of looming and compound trajectories, compared to translating trajectories (Friedman Repeated Measures Analysis of Variance on Ranks, χ^2^ = 440.996, *p* < 0.001) (Fig 3.d). Furthermore, units responded to a larger proportion of direct looms (Loom), compared to Compound A-P.

We observed similar trends when pooling all units based on stimulus type. The number of responsive units was compared among different trajectories and velocities. As shown in Fig 4, the stimulus trajectory significantly influenced the number of responsive units (Two-way RM ANOVA, F_4_ = 98.781, *p* < 0.001). Specifically, trajectories involving direct looms or compound trajectories with a looming phase evoked a greater number of responsive units, compared to translating trajectories (Holm-Sidak post-hoc test, *p* < 0.001). Consistent with the findings from individual units, Loom evoked a higher number of responsive units compared to Compound A-P, while no difference was observed between the two compound trajectories, Compound A-P and Compound P-A (Holm-Sidak post-hoc test, t = 2.871, *p* = 0.02), and no difference was found between the two translating trajectories, Translation P-A and Translation A-P (Holm-Sidak post-hoc test, t = 1.355, *p* = 0.327). Additionally, the variance in the number of units responding to Compound A-P was larger across different velocities, with the slowest velocity (V5, *l*/|*v*| = 50 ms) eliciting fewer unit responses compared to Compound P-A and Loom. This discrepancy largely contributed to the difference between Loom and Compound A-P.

**Figure 4.**
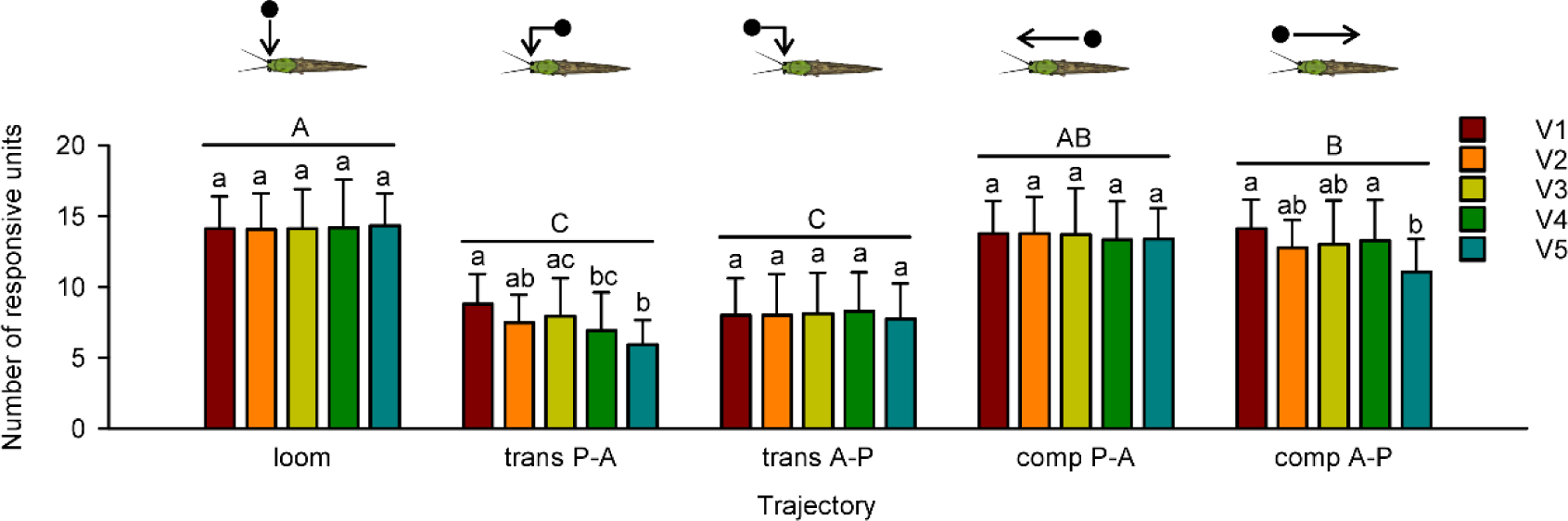
Comparison of the number of units that responded to each visual stimulus. Upper case letters represent the statistical comparisons between different stimulus trajectories. Lower case letters represent the comparison between different velocities (converted to *l*/|*v*|) within each stimulus category.

### Response categories

Responsive units were pooled across all animals for each stimulus type, and their peristimulus time histograms (PSTHs) were visually inspected and categorized based on the pattern of firing rate changes over time. Specifically, for Loom at V1 (*l*/|*v*| = 11.7 ms), five categories of responses were identified (Fig 5).

**Figure 5.**
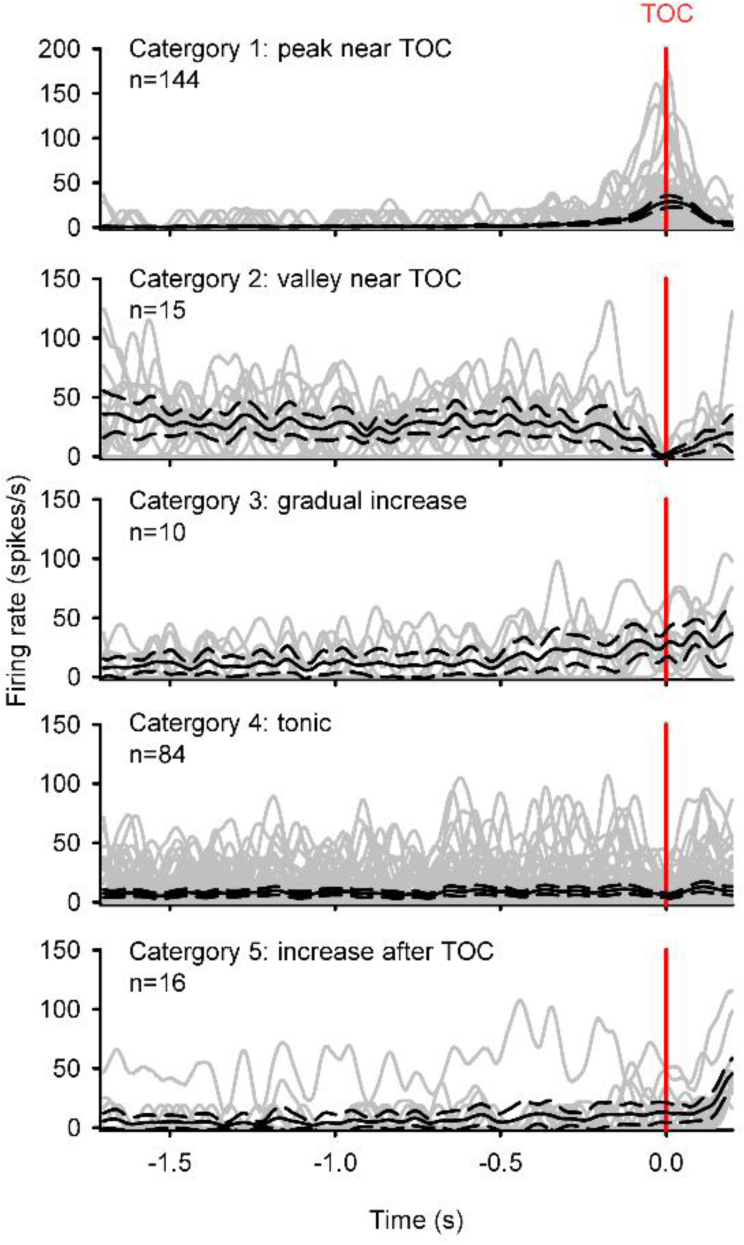
Categorization of responsive units when the animal was presented with a looming stimulus (*l*/|*v*| = 11.7 ms). All peristimulus time histograms (PSTHs), represented by the gray lines, were aligned to the projected time of collision (TOC), represented by the red vertical line. The black lines represent the average PSTH for the plot. Category 1 included units that showed a peak near TOC. Category 2 included units that showed a valley near TOC. Category 3 included units that showed a gradual increase during the stimulus presentation. Category 4 included units that showed tonic firing at a relatively steady rate during the stimulus presentation. Category 5 included units that showed an increase after TOC.

Category 1 comprised units that exhibited a peak firing rate near the projected time of collision (TOC). Category 2 included units that showed a valley in firing rate near TOC. Category 3 consisted of units that displayed a gradual increase in firing rate over time. Category 4 comprised units that maintained a relatively consistent firing rate in a tonic manner. The last category included units that displayed a tonic firing rate before TOC, with an increase in firing rate near the end of the recording. It is worth noting that since the setup was open-loop, and the visual stimulus stopped expanding 0.08 seconds before TOC, the firing rate increase observed at the end was likely unrelated to the looming stimulus.

Among the 269 units that responded to a Loom at V1, 144 units fell into category 1 (53.5%), 15 units fell into category 2 (5.6%), 10 units fell into category 3 (3.7%), 84 units fell into category 4 (31.2%), and 16 units fell into category 5 (5.9%). Excluding categories 4 and 5, which consisted of units that did not exhibit clear changes in firing rate over time, the majority of units responded to Loom with a peak near TOC. This pattern of response is consistent with the known response characteristics of DCMD and LDCMD neurons. Other visual stimulus trajectories revealed similar response categories, with the majority of units displaying DCMD-like response patterns. For a more objective comparison, we extracted common trends that statistically represent the response category and corresponding distribution.

### Common Trends Responding to Looming Stimuli

Dynamic factor analysis (DFA) was performed using the MARSS package in R 4.1.3 to extract common trends (CTs) from the responsive units. However, due to the computational complexity of performing DFA on Gaussian-smoothed 1-ms-binned data, which consistently caused R to crash, an alternative approach was employed. The firing rate was recalculated using a 50-ms bin width without smoothing. Although this reduced the temporal resolution, it still captured the dynamic modulation of the firing rate while significantly reducing computational demands by a factor of 50.

For each of the 25 visual stimuli, DFA was performed iteratively with an increasing number of common trends, starting from 1. The Akaike information criterion with a correction for small sample sizes (AICc) was used to monitor the quality of each model. The iteration stopped when the AICc started to increase, and the model with the lowest AICc was selected as the optimal model. Supplementary Table 3 summarizes the optimal number of common trends for looming, translating, and compound trajectories, which were 7, 3, and 5, respectively. From the DFA model, the *i-th* column of the hidden process matrix (**x**) represents the instantaneous firing rate of the *i-th* common trend. The peristimulus time histograms (PSTHs) of the common trends were plotted using SigmaPlot and aligned using the projected time of collision (TOC) or the time that the object passed 90°azimuth (T90) (Fig 6).

**Figure 6.**
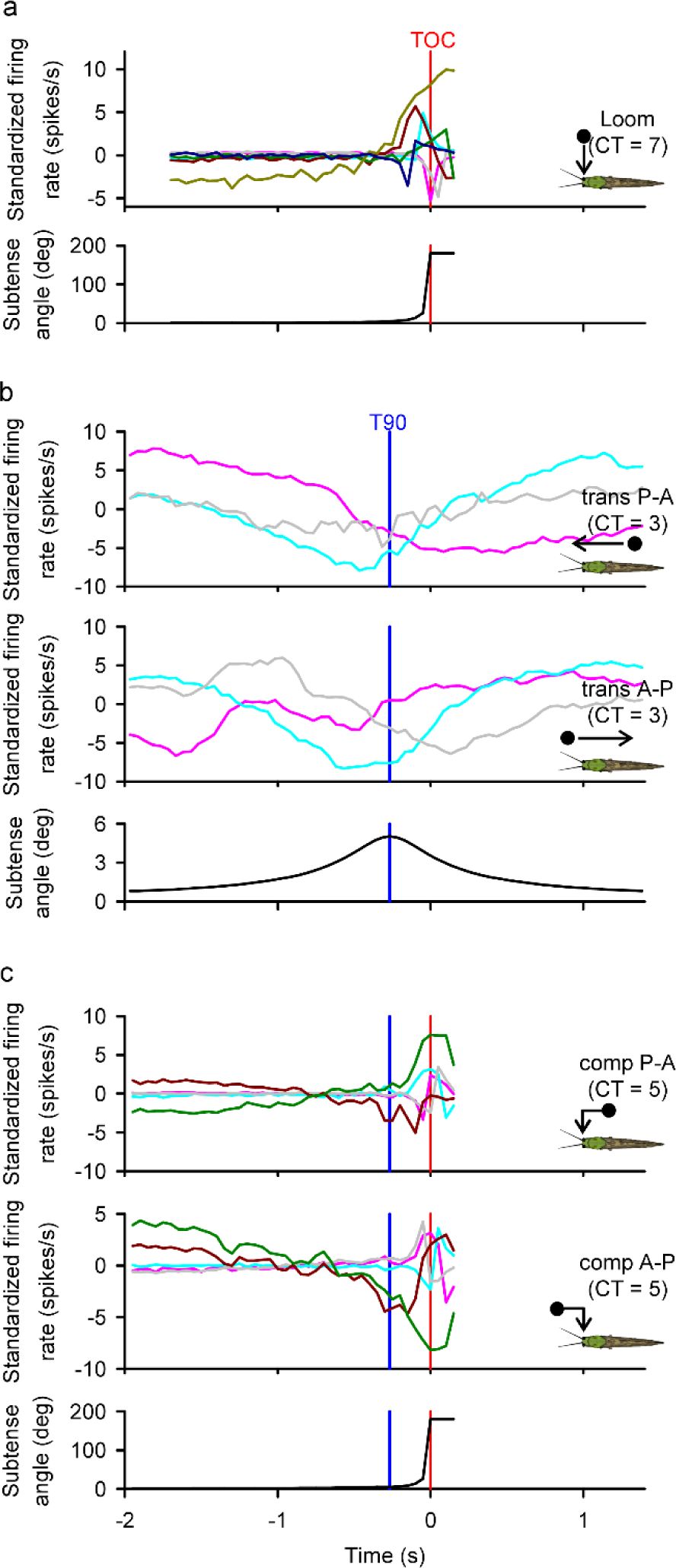
Peristimulus time histograms of population responses (common trends) from the best DFA models for each of the 5 trajectories at V1 (*l*/|*v*| = 11.7 ms), and the real-time subtense angle of the stimulus. The common trends were aligned to either projected time of collision (TOC, red vertical line) or the time of 90 degrees azimuth (T90, blue vertical line). Note that for the translational trajectories, the maximum subtense angle was ∼5 degrees, and thus the y-axis was scaled differently. “CT” refers to the number of common trends that describe the response for each trajectory.

Fig 7 illustrates the PSTHs of the 7 common trends in response to direct looms at different velocities (V1-V5), and the corresponding factor loadings of units in each common trend as heat maps. Notably, the order of common trends in the DFA model is randomized, making direct comparisons between the response of each common trend across different visual stimuli impossible. Despite the various types of unit responses categorized in Fig 5, 6 out of 7 common trends exhibited a clear firing rate peak, resembling Category 1 responses from discriminated units. The remaining trend at each velocity (CT6 at V1, V2, and V3, CT7 at V4 and V5) displayed a consistently increasing or decreasing firing rate similar to Category 3 responses. While most common trends showed positive peaks, some common trends, such as CT5 at V2 and CT4 at V3, displayed negative peaks. As stated in the Materials and Methods, the common trends have been re-scaled and reversed based on the average factor loadings. However, in the common trends with negative peaks, the largest values in the factor loadings (represented by the bars with the darkest colour) were mostly negative (blue). This suggests that although the plotted common trends contribute positively to all units on average, the top units that these negatively peaking common trends contributed to, i.e., the units that correlated best with these common trends, actually have positive peaks. Correspondingly, it is worth noting that all units in Response Category 1 showed positive peaks (Fig 4). Although Response Category 2 units showed valleys near TOC, the number of units in Category 2, as well as the amplitude of the valleys, were much lower than in Category 1. In summary, the dominant type of response among the common trends was peaking positively near TOC. Therefore, in subsequent analysis, the common trends with negative peaks were flipped vertically to accurately represent the responses of the majority of units.

**Figure 7.**
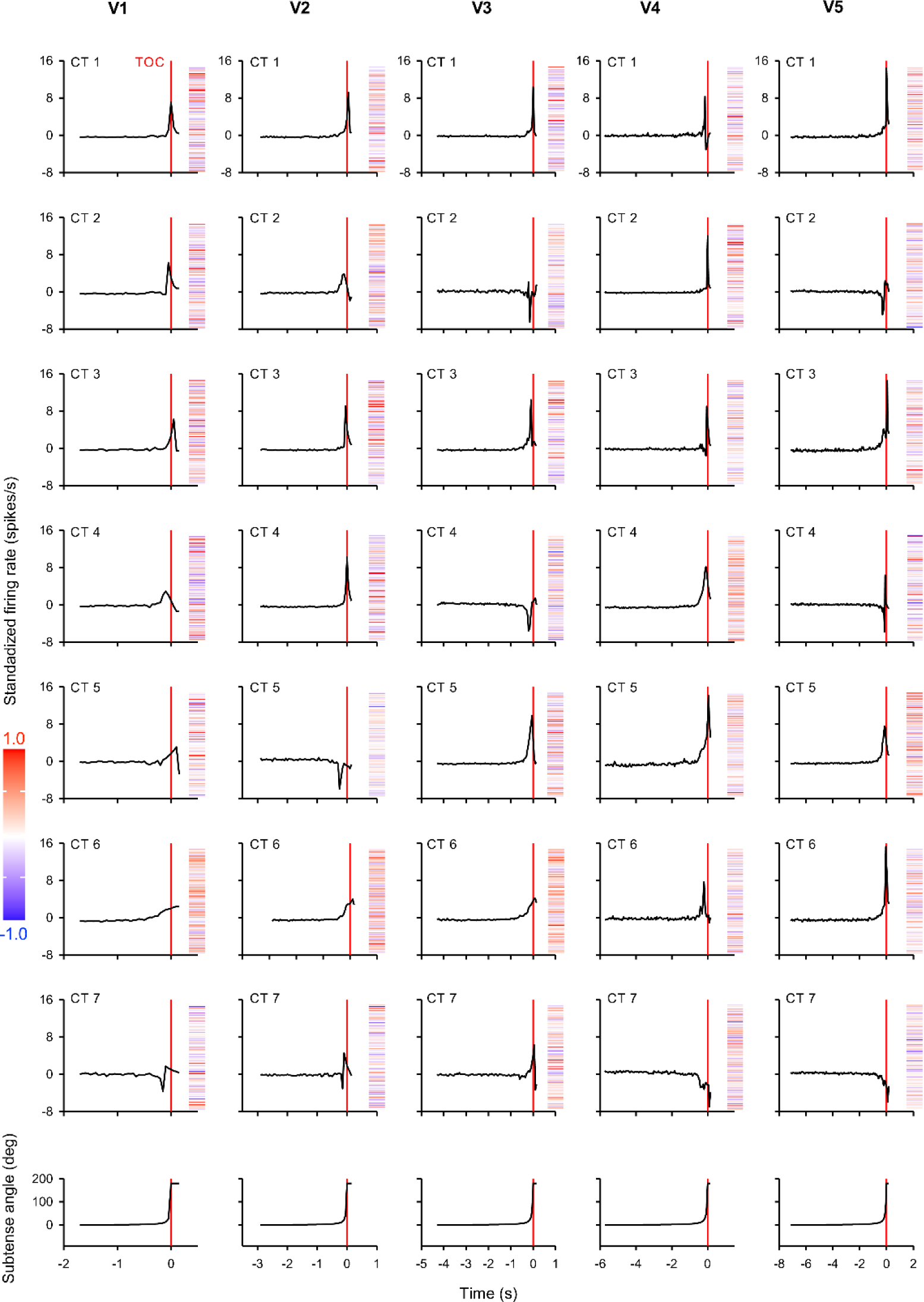
Common trends (CTs) evoked by direct looming stimuli at 5 different velocities (*l*/|*v*|). For looming stimuli, the best DFA model was determined to contain 7 CTs. In each column (approach velocity), peristimulus time histograms (PSTHs) of the 7 CTs were plotted, aligned to projected time of collision (TOC). In DFA models, the order of CTs was randomized, i.e., CT1 from the first and second column are not necessarily the same. The factor loadings of each unit (total = 411 from 19 locusts) for the CTs are shown next to the PSTH in a heatmap in which the loadings ranged from +1 (red) to -1 (blue, see Materials and Methods for details). Most CTs showed a positive peak near TOC, while some units (e.g. contributing to CT 5 at V2) showed a negative peak.

From the common trends exhibiting a clear peak, various parameters were measured and compared between different velocities, including peak time, subtense angle at peak (peak SA), half peak time (when the firing rate reached 50% of the peak firing rate), subtense angle at half peak (half peak SA), peak width at half height (PWHH), half-peak-to-peak time, and peak-to-half-peak time (Fig 8). Slower velocities evoked earlier peak times relative to TOC. However, the peak SA increased with slower velocities, indicating that the peak occurred when the object was closer to the locust. The half peak time and half peak SA displayed similar trends: slower velocities caused the half peak to occur earlier, but the half peak SA increased, meaning that half peak also occurred when the object was closer. Both the half-peak-to-peak time and peak-to-half-peak time increased as the stimulus velocity decreased, contributing to an overall increase in PWHH.

**Figure 8.**
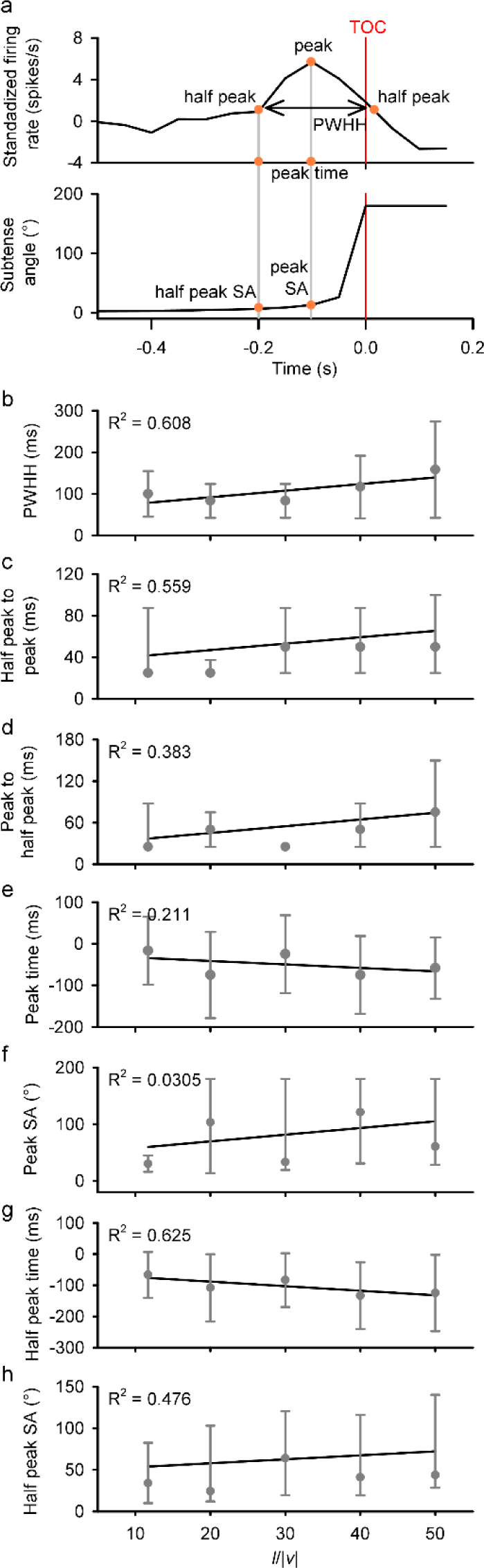
Response parameters of the common trends that peaked near TOC. The peak time, half peak time (when firing rate reached half of the peak firing rate), peak width at half-height (PWHH), half-peak to peak time, peak to half-peak time, and the subtense angle (SA) at peak and half peak were compared between different values of *l*/|*v*|. a) A sample PSTH of a common trend and the subtense angle of the corresponding visual stimulus, with yellow circles highlighting the peak and half peak time, the peak and half peak subtense angle (SA), and the peak width at half height (PWHH). b) As *l*/|*v*| increased, both half-peak-to-peak and peak-to-half-peak increased, resulting an increased PWHH. c) As *l*/|*v*| increased, However, the SA of the stimulus at the peak increased with l/|v|. d) As *l*/|*v*| increased, the half peak also occurred earlier, indicating that the responses started earlier. The SA of the stimulus at half peak increased with *l*/|*v*|. It is worth noting that none of the linear regressions were statistically significant. This is likely due to the reduced resolution when 50-ms bin width was used for the DFA models. Parametric parameters (b, e, g) were plotted as mean ±standard deviation, while non-parametric parameters (c, d, f, h) were plotted as median with positive and negative error bars as the 75^th^ and 25^th^ percentiles, respectively. A linear regression was fitted between parameter averages and *l*/|*v*|.

Although none of the comparisons yielded statistically significant results (Peak time: One-way ANOVA, F_4_ = 0.568, *p* = 0.688; Peak SA: Kruskal-Wallis One Way Analysis of Variance on Ranks, H_4_ = 1.757, *p* = 0.780; Half peak time: One-way ANOVA, F_4_ = 0.462, *p* = 0.763; Half peak SA: Kruskal-Wallis One Way Analysis of Variance on Ranks, H_4_ = 1.995, *p* = 0.737; PWHH: One-way ANOVA, F_4_ = 1.148, *p* = 0.357; Half-peak-to-peak time: Kruskal-Wallis One Way Analysis of Variance on Ranks, H_4_ = 2.261, *p* = 0.688; Peak-to-half-peak time: Kruskal-Wallis One Way Analysis of Variance on Ranks, H_4_ = 6.294, *p* = 0.178), it is important to note that the power of all these tests was below the desired power of 0.8. This lack of statistical significance can be attributed to the reduced temporal resolution (50-ms bin width) and the relatively small sample size (n=6).

In summary, the majority of discriminated units respond to looms with a peak firing rate near TOC, irrespective of the looming velocity. Similar to the response of LGMD/DCMD, the peak time and half peak time of the common trends occurred earlier with slower looming velocities, however the subtense angle at peak and half peak firing rate were both higher.

### Common Trends Responding to Translations

In response to translational movement, a DFA model with three common trends was determined to be the most optimal (Supplementary Table 3, Fig 9). Consistent with the characteristics of the LGMD/DCMD pathway, discriminated responsive units displayed a lower sensitivity to translations, while extracted common trends exhibited more variation in their peristimulus time histograms (PSTHs). Notably, translational movements initiated from the anterior visual field (Translation A-P) or the posterior visual field (Translation P-A) evoked common trends with similar PSTHs.

**Figure.9.**
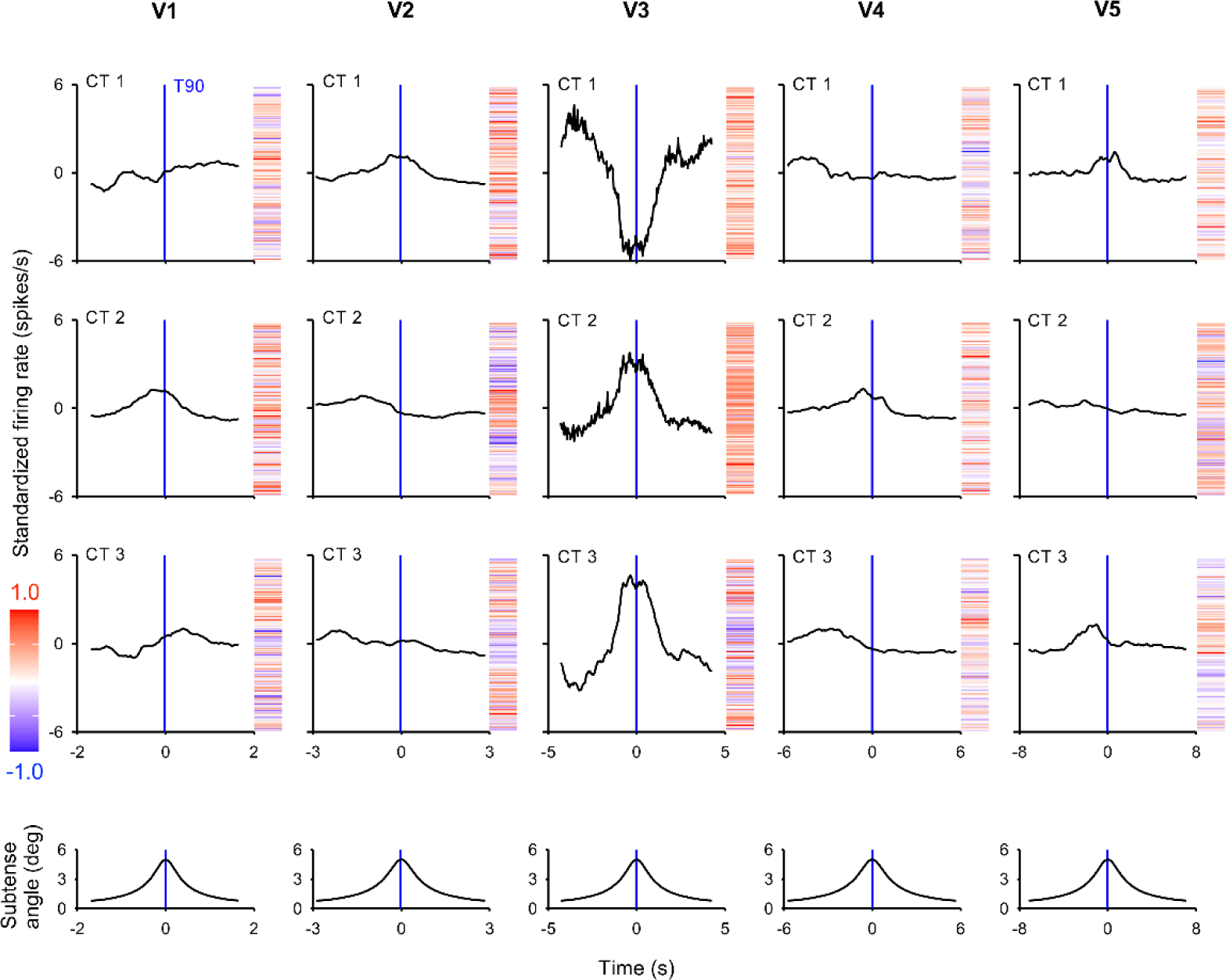
Common trends (CTs) of translational stimulus (anterior to posterior) at 5 different *l*/|*v*|. For translational stimuli, the best DFA model was determined to contain 3 CTs. In each column, peristimulus time histogram (PSTH) of the 3 CTs were plotted, aligned to the time at 90 degrees azimuth (T90). The factor loadings of each CT are shown next to the PSTH in a heatmap. At least one CT at each velocity peaked near T90, followed by a gradual decay, similar to the response of the DCMD. Other CTs maintained a relatively stable firing rate without clear change near T90.

Among the 15 CTs observed in response to Translation A-P, 7 CTs (CT2 at V1, CT1 at V2, CT2&3 at V3, CT 2 at V4, and CT 1 at V5) displayed a gradual increase in firing rate, peaked near the time of 90°azimuth (T90), and gradually decreased after the stimulus passed T90, resembling the response characteristics of DCMD. Other response patterns included an early peak (e.g., CT3 at V2) or a gradual increase without clear tuning near T90 (e.g., CT1 at V1). In response to Translation P-A, 6 out of 15 CTs peaked near T90, while others displayed the alternative response types observed in Translation A-P. These response patterns were also observed in the response to the translating phase of the compound trajectories.

### Common Trends Responding to Compound Stimuli

Fig 10 presents the common trends observed in response to the compound trajectory (Compound A-P) at five different velocities. As described above, common trends with predominantly negative factor loadings were inverted to accurately reflect the majority of unit activity, and the factor loadings were scaled from -1 to 1. The peristimulus time histograms (PSTHs) were aligned to both the time the object passed through 90°azimuth (T90) and the projected time of collision (TOC). Similar to the responses to looming stimuli, most of the common trends in response to the compound trajectories exhibited a distinct peak near TOC, although CT5 at V1 showed a continuous increase until the end of the trial. However, the PSTHs of the peaking common trends were also influenced by the translating phase of the trajectory.

**Figure.10.**
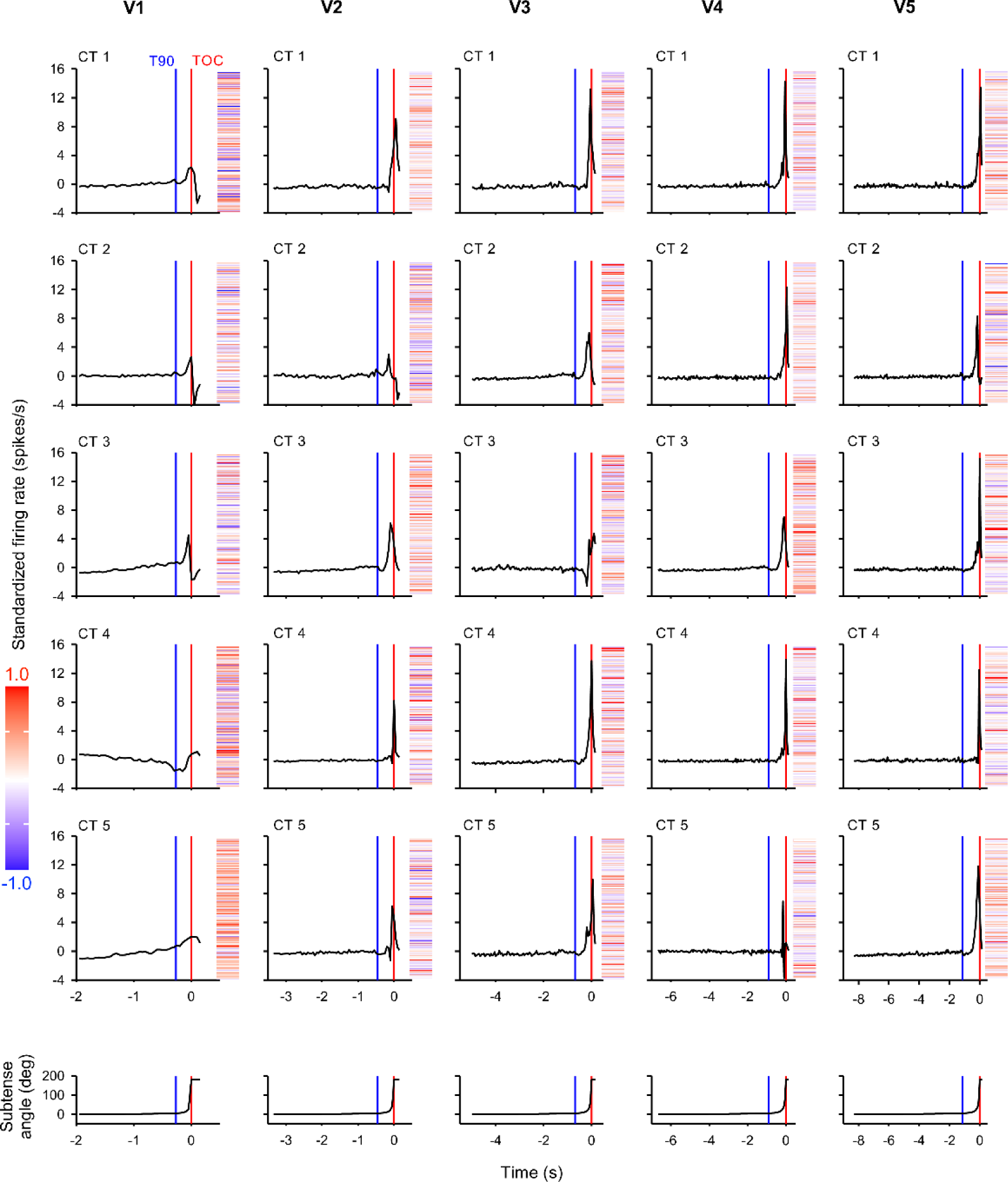
Common trends (CTs) of compound stimulus (anterior to posterior) at 5 different *l*/|*v*|. For compound stimuli, the best DFA model was determined to contain 5 CTs. In each column, peristimulus time histogram (PSTH) of the 5 CTs were plotted, aligned to both projected time of collision (TOC) and time at 90 degrees azimuth (T90). The factor loadings of each CT are shown next to the PSTH in a heatmap. At V1, CT1 and 2 both displayed a gradual increase during the translational phase, a small peak near T90, a brief valley after T90, and then a larger peak near TOC, similar to the response of DCMD. CT3 displayed gradual increase before peaking near TOC. CT 4 firing rate first decreased during the translating phase, and then increased towards the end of the stimulus. CT 5 firing rate gradually increased towards the end of stimulus presentation. As velocity decreased, the variation among CTs decreased. At V2 and V3, 4 out of 5 CTs displayed a constant low firing rate before peaking near TOC, while at V4 and V5, all five CTs displayed this type of response.

At V1, CT1 and 2 displayed a small peak near T90, followed by a valley, and then a larger peak near TOC. This response pattern is consistent with the activity of DCMD in response to a similar motion trajectory observed in previous studies (McMillan and Gray 2012; Dick and Gray 2014). During the translating phase of the trajectory, CT3 at V1 exhibited a gradual increase in firing rate, followed by a rapid rise to a peak around TOC during the looming phase. This gradual increase in firing rate was also observed in response to pure translational movement (Fig 9). CT4 at V1 displayed a valley near T90, followed by a continuous increase until after TOC, suggesting that a subgroup of units was inhibited by the translating stimulus but responsive to looming.

The variation in CT responses decreased as the stimulus velocity decreased. At V2 and V3, only one unit displayed alternative response: CT2 at V2 was similar to CT1 at V1, while CT3 at V3 displayed a large valley before increasing towards TOC. All other four CTs maintained a tonic firing rate before peaking near TOC, similar to the response to simple looms. At V4 and V5, all CTs exhibited similar response patterns. The distinct valley after T90, observed in the DCMD response to compound trajectories, becomes less pronounced as the stimulus velocity decreases (Dick and Gray 2014). This was also observed among the common trends derived from all discriminated units reported here.

## Discussion

Previous studies have quantitatively described the response of individual neurons, such as the LGMD/DCMD pathway, to visual stimuli along different trajectories at various velocities (Rind and Simmons 1992; Simmons and Rind 1992; McMillan and Gray 2012; Dick and Gray 2014; Stott et al. 2018). However, the response of other motion-sensitive neurons to object motion along complex trajectories at different velocities has not been described on a population level. We showed here that multiple units (neurons) respond to object motion along looming, translating, or compound trajectories that transitioned from translation to loom. Common trends (CTs), which represent the activity of neural ensembles, were extracted using the dynamic factor analysis (DFA) and compared between different trajectories and velocities. Based on the model selection criteria, there were 7 CTs among units responding to looms, 3 CTs among units responding to translations, and 5 CTs among units responding to compound trajectories.

### Tetrode recording

The custom-fabricated tetrode we used here was inserted parallel into the ventral nerve cord, and surrounded by the sheath, which, combined with the hook electrode, provided a stable support of the nerve cord during recording. The preparation required manual lifting of the ventral nerve cord and insertion of the tetrode. In the connective between the suboesophageal and prothoracic ganglion, there are ∼3000 neuron axons (Rowell and Dorey 1967), and we only recorded from a small subset (mean = 23, S.D. = 2) of these neurons in each animal. Since the positioning of the tetrode was not identical in each preparation, it was not possible to record from the same neurons every time. However, Fig 3.c shows that the number of stimuli individual units responded to was consistent across all 19 animals, indicating that despite slight positioning differences, the units that were recorded in different animals were generally from the same population. Furthermore, the slight difference in the positioning of tetrodes can, in fact, help increase the diversity of units being recorded. Across 19 animals, this will increase the percentage of descending interneurons being sampled. It was also possible that the same unit was recorded repeatedly in different animals. This was solved by pooling units from all animals and performing dimensional reduction procedures, such as the dynamic factor analysis (DFA). The same unit was expected to respond similarly between different animals, and thus would be represented by the same hidden process (common trend).

### Dynamic factor analysis

Large number of neurons that respond to a given stimulus usually display lower-dimension activities, and the coupling between neighbouring neurons is highly diverse, contributing to the diverse population dynamics (Okun et al. 2015). Therefore, we performed dynamic factor analysis (DFA) to extract common trends in response to different types of visual stimuli. We plotted the peristimulus time histograms (PSTHs) of individual unit responses with 1-ms bin and 50-ms Gaussian smoothing window. However, when performing DFA, due to the limitation of computational power of the R package and the computing devices that we could access, we down-sampled the data with 50-ms bins. Although this resulted in a reduced temporal resolution, similar binning widths were used in earlier studies (30 ms: (Pinter et al. 1982); 25 ms: (Simmons and Rind 1992); 12.5 ms: (Gray et al. 2001); 10 ms: (Rind and Simmons 1997; Santer et al. 2008)) and they were sufficient to describe the dynamic firing rate change of LGMD/DCMD. More recently, 50-ms bin was used in a similar analysis of population coding of visual information in locusts (Parkinson et al. 2020). Moreover, the contribution of DCMD in the generation of avoidance behaviour is dependent on the wing beat cycle, which is around 50 ms per cycle (Santer et al. 2006), suggesting that shorter bin width is not necessarily more biologically relevant. Therefore, the longer bin width is still capable of describing the activity of motion-sensitive neural ensembles in response to various motion trajectories.

### Response of neuron population based on stimulus types

*Looming*. Previously identified motion-sensitive descending interneurons, such as LGMD/DCMD and the LDCMD, respond to looms with increased firing rate that peaks near the projected time of collision (TOC), and looming stimuli approaching from 90°azimuth evokes the highest response (Rogers et al. 2010). Among all discriminated units that responded to looms, we found that more than half (53.5%) of the units responded with a peak near TOC, similar to the response of the LGMD/DCMD pathway. This was further confirmed by the distribution of extracted common trends, where 6 out of 7 common trends peaked near TOC. The last CT shows a constant increase towards the end of the trials. The remaining units either responded with a valley near TOC or no clear peak/valley, neither of which were reflected in the common trends, indicating that these units contribute little to the population coding. Despite no statistical significance, timings of the CTs in response to looming at different velocities displayed a similar trend to DCMD (Dick and Gray 2014), suggesting a universal encoding strategy among all looming-sensitive neurons.

*Translation.* Compared to looms, translations, which are not on a collision course, usually represent less danger. In locusts, the LGMD/DCMD pathway is less sensitive to translating stimuli, compared to looming (Rind and Simmons 1992; Hatsopoulos et al. 1995; Gabbiani and Krapp 2006; Peron and Gabbiani 2009; McMillan and Gray 2012). We found that translating stimuli evoked fewer units to respond, and the number of CTs (n=3) was less than that evoked by looms (n=7), indicating less variation across the unit responses. This is biologically relevant, since approaching objects often represent predators or possible collisions, and thus require more attention. Small target motion detectors that show little or no response to larger objects were identified in cats (Hubel and Wiesel 1965), dragonflies (O’Carroll 1993), hoverflies (Nordström and O’Carroll 2006), and fruit flies (Keleş and Frye 2017). In fruit fly *Drosophila*, the lobula columnar (LC) neurons aggregate retinotopic inputs and project onto different optic glomeruli in the protocerebrum (Strausfeld and Okamura 2007). Some types of LC neurons are sensitive to looming stimuli, while others are sensitive to small objects that are not on a looming course (Wu et al. 2016). To our knowledge, neurons that respond exclusively to non-looming objects have not been identified in locusts (Kien 1974; Borst and Haag 2002).

It is also worth noting that the definition of translation is not consistent in previous studies. For example, (Peron and Gabbiani 2009) defined translation as a trajectory that maintains the same distance from the locust, which was a rotary movement around the locust. In this study, the translating stimuli travelled along a straight trajectory parallel to the longitudinal axis of the animal, meaning that the subtense angle increased as the stimulus approached 90°azimuth. Therefore, it contained a looming element in addition to the rotary trajectory used in (Peron and Gabbiani 2009). In other studies, such as (Judge and Rind 1997), the “near-miss” trajectory was in fact the same as the translating trajectory used here. As the trajectory deviated further from a direct collision course, DCMD response decreased, in both the number of spikes and the peak firing rate (Judge and Rind 1997).

For the translating trajectories, as well as the compound trajectories, the distance between the disc and the locust was set at 80 cm. This distance was used in previous studies on DCMD (McMillan and Gray 2012; Dick et al. 2017). When passing through 90°azimuth (T90), the subtense angle of the disc was 5.01°. In tethered flying locusts, near-miss looming stimuli that have a maximal subtense angle of 10°evoked avoidance behaviours (Robertson and Johnson 1993). The 80 cm distance was chosen to further differentiate the translations from “near-miss” looms.

*Translation transitioning to loom.* When a predator, such as a bird, spots a locust and turns towards it, the trajectory of the bird will be compound, initiating as a translation and transitions to a loom. Since the discriminated neuron population did not respond much to translating stimuli, it is interesting to study the response to a compound trajectory. In response to a disc moving along a compound trajectory, the number of responsive units was similar to that responding to a simple loom, however the number of CTs among responsive units (n=5) was fewer than that evoked by looms (n=7) but more than simple translations (n=3). DCMD responds to a similar compound trajectory with a gradual increase, followed by a valley right after the time of transition (TOT), similar to T90, and a higher peak near TOC (McMillan and Gray 2012; Dick and Gray 2014), which can be observed in CT1 at V1, but not other CTs. Most CTs look similar to those evoked by a simple loom. Overall, the neuron ensembleis capable of responding to compound trajectories with similar magnitude as looms, although the looming phase is relatively short, and the variation was less.

### Directional selectivity of descending motion-sensitive interneurons

DCMD responds to translating visual stimuli with an increasing firing rate that peaks near T90 (McMillan and Gray 2012). When comparing translation that initiated from the anterior (A-P) to translation that initiated from the posterior (P-A), Translation A-P causes a more acute increase in DCMD firing rate and a higher peak in DCMD response (McMillan and Gray 2012). We found that discriminated units were more responsive to Translation A-P: out of the five velocities in each translating trajectory, the discriminated units responded to more types of Translation A-P (median = 1, 1^st^ quartile = 0, 3^rd^ quartile = 3), compared to Translation P-A (median = 1, 1^st^ quartile = 0, 3^rd^ quartile = 2), and as a result, the number of units that responded to Translation A-P was higher than Translation P-A, although neither comparison was statistically significant. The LGMD/DCMD pathway is more sensitive to translating motion initiated from the anterior side, compared to the posterior side (Peron et al. 2009; Rogers et al. 2010). We found that this trend might be common among other motion-sensitive neurons as well. However, it is worth noting that this trend is reversed for compound trajectories: more units responded to Compound P-A, compared to A-P. Moreover, the effect of motion velocity was larger for Compound A-P, compared to Compound P-A. Using the current approach, it is difficult to distinguish which phase of the compound trajectory the units responded to. This difference could be a result of the transition from translation to loom.

As discussed above, directional small-field motion-sensitive neurons have been identified in several species, such as cats (Wickelgren and Sterling 1969), crabs (Scarano et al. 2020), and hawkmoth (Collett 1971). However, in locusts, the only directionally selective neurons identified thus far respond to large-field movements, but not small translating objects (Rind 1990a, b). In the locust compound eye, the ommatidia density is higher in the anterior side, compared to the posterior side (Krapp and Gabbiani 2005; Peron et al. 2007), however the LGMD dendrites sampling the posterior visual field are thicker (Peron et al. 2007), resulting in DCMD responding more to objects looming within the posterior visual field (Krapp and Gabbiani 2005). Overall, the population of motion-sensitive neurons did not display statistically significant preference to motion towards a specific direction. However, our results could not exclude the possibility that some units respond differently to motion towards different directions, and the responses of such units were masked by the activity of other units in the common trends. Although these units did not contribute significantly to the common trends, specific synaptic projections from these units could still play an important role in generating appropriate behavioural responses. Further studies, possibly involving intracellular recordings, would be necessary to investigate the response preference of individual units and their role in controlling avoidance behaviours.

### Dynamic composition of neural ensembles

Despite variations in brain sizes among different animals, all rely on a finite number of neurons to execute a wide range of tasks. In the late 1940s, it was proposed that individual neurons could be part of different neural assemblies depending on functional requirements (Hebb 1949). For example, in mouse brain, memories are stored in a sparse population of neurons called engram cells (Josselyn and Tonegawa 2020). Although traditional theories assume engrams to be stable, in order to ensure stable behavioural outputs, recent studies have found that the composition of engrams is, in fact, fluid and flexible (Sweis et al. 2021). The contribution of individual neurons to an engram reconfigures over periods of weeks, likely contributing to memory flexibility yet ensuring the population output remains largely consistent (Driscoll et al. 2017; Rule et al. 2020).

In sensory processing, a large number of neurons respond to a single sensory modality, while their activity can be represented by a lower dimension, via the dynamic correlation of individual neurons (Kenet et al. 2006). In the mouse visual cortex, neighbouring neurons display diverse levels of coupling with the population coding (Okun et al. 2015). Although the strength of synaptic connections fundamentally determines the level of coupling, and thus is independent of response preferences or stimulus amplitude (Jacobs et al. 2015), it can be affected by non-sensory variables like motor intention (Okun et al. 2015). The dimensionality of ensembles in the rat gustatory cortex can also grow with ensemble size (Mazzucato et al. 2016).

In smaller nervous systems with fewer neuron counts, it is even more important to allow dynamic composition of neural ensembles, in order to fulfill the computational needs. In the moth antennal lobe, groups of units coordinate to form unique population-level responses to various olfactory stimuli (Daly et al. 2004). In the cockroach central complex, multiple units that are sensitive to mechanical stimulation also respond to changes in velocity and light, and their population activity likely governs corresponding behavioural responses (Ritzmann et al. 2008). In locusts, the population response of motion-sensitive neurons is affected by the treatment of sublethal doses of pesticides, corresponding to other physiological and behavioural changes (Parkinson et al. 2020).

We found that although the responsive units generally came from the same population, the dimensionality of the ensembles was affected by the stimulus trajectory (Figure 6). The contribution of individual units to different ensembles is also dynamic, for the same motion trajectory presented at different velocities (Figure 7). Although the correlation between individual units and each common trend was not directly assessed here, due to the pooling of units from all animals, the common trend dynamics can still reflect the flexibility of ensemble configuration. Since the population coding, instead of individual neuronal activity, is recognized by downstream motorneurons to guide behavioural responses, reconfiguration of neural ensembles can provide a more detailed and comprehensive description of the external environment.

## Author contributions

Conceptualization: Sinan Zhang and John Gray; Methodology: Sinan Zhang and John Gray; Formal analysis and investigation: Sinan Zhang; Writing - original draft preparation: Sinan Zhang; Writing - review and editing: Sinan Zhang and John Gray; Funding acquisition: John Gray; Resources: John Gray; Supervision: John Gray.

## Statement and Declarations

Funding was provided by the Natural Sciences and Engineering Research Council of Canada (Award RGPIN-2019-03983). The authors declare no conflicts of interest, financial or proprietary.

