## Supplementary files for "Complex motion trajectories are represented by a population code from the ensemble activity of multiple motion-sensitive descending interneurons in locusts"

**Supplementary Table 1** Summary of spike detection and unit discrimination. On average, we detected 41528 spikes (waveforms), and discriminated 23 units from each animal. The MANOVA results show that the discriminated units were statistically distinct from each other.

| Animal | Total Number of Spikes | Number of Units | MANOVA (3D space) |
|---|---|---|---|
| Locust 01 | 35428 | 22 | $F(63,105681) = 23.144279$, $p < 0.001$ |
| Locust 02 | 42581 | 22 | $F(63,119487) = 18.639135$, $p < 0.001$ |
| Locust 03 | 53311 | 26 | $F(75,139210) = 14.34502$, $p < 0.001$ |
| Locust 04 | 63911 | 25 | $F(72,169609) = 14.117809$, $p < 0.001$ |
| Locust 05 | 52634 | 24 | $F(69,156198) = 28.1962$, $p < 0.001$ |
| Locust 06 | 39845 | 22 | $F(63,113609) = 24.221551$, $p < 0.001$ |
| Locust 07 | 41466 | 25 | $F(72,116124) = 9.952913$, $p < 0.001$ |
| Locust 08 | 33611 | 21 | $F(60,95534) = 30.451603$, $p < 0.001$ |
| Locust 09 | 50010 | 23 | $F(66,142358) = 43.587911$, $p < 0.001$ |
| Locust 10 | 53630 | 20 | $F(57,129504) = 16.581168$, $p < 0.001$ |
| Locust 11 | 35743 | 24 | $F(69,105748) = 29.223666$, $p < 0.001$ |
| Locust 12 | 31739 | 25 | $F(72,94772) = 35.440023$, $p < 0.001$ |
| Locust 13 | 50987 | 24 | $F(69,142730) = 27.030042$, $p < 0.001$ |
| Locust 14 | 28959 | 22 | $F(63,86371) = 20.296337$, $p < 0.001$ |
| Locust 15 | 27028 | 22 | $F(63,80607) = 57.74843$, $p < 0.001$ |
| Locust 16 | 33660 | 20 | $F(57,100299) = 25.223018$, $p < 0.001$ |
| Locust 17 | 29110 | 23 | $F(66,86858) = 20.473046$, $p < 0.001$ |
| Locust 18 | 53089 | 24 | $F(69,140337) = 41.09711$, $p < 0.001$ |
| Locust 19 | 32286 | 22 | $F(63,88389) = 56.060543$, $p < 0.001$ |
| **Total** | 789028 | 436 | |
| **mean** | 41528 | 23 | |
| **median** | 39845 | 23 | |
| **s.d.** | 10821 | 2 | |
| **min** | 27028 | 20 | |
| **max** | 63911 | 26 | |
| **1st quartile** | 32286 | 22 | |
| **3rd quartile** | 52634 | 24 | |

**Supplementary Table.2** Response of all discriminated units from one animal to all 25 visual stimuli. An orange block indicates that the unit responded to this specific visual stimulus, while a blue box indicates that this unit did not respond.

| Stimulus | Discriminated units | | | | | | | | | | | | | | | | | | | | | |
|---|---|---|---|---|---|---|---|---|---|---|---|---|---|---|---|---|---|---|---|---|---|---|
| | a | b | c | d | e | f | g | h | i | j | k | l | m | n | o | p | q | r | s | t | u | v |
| loom V1 | orange | blue | orange | blue | blue | blue | orange | blue | blue | orange | orange | blue | orange | blue | blue | orange | orange | orange | orange | orange | blue | blue |
| loom V2 | blue | orange | orange | blue | blue | orange | orange | orange | orange | orange | orange | orange | orange | blue | blue | orange | orange | orange | orange | orange | orange | blue |
| loom V3 | orange | orange | orange | orange | orange | orange | orange | blue | orange | orange | orange | blue | orange | blue | blue | orange | orange | orange | blue | orange | orange | blue |
| loom V4 | blue | blue | orange | blue | blue | orange | orange | blue | blue | orange | orange | blue | orange | blue | blue | orange | orange | blue | blue | orange | orange | blue |
| loom V5 | orange | orange | orange | orange | blue | orange | orange | orange | orange | orange | orange | blue | orange | blue | orange | blue | orange | blue | blue | orange | blue | blue |
| trans P-A V1 | orange | orange | orange | blue | blue | blue | blue | orange | blue | blue | orange | blue | blue | blue | blue | blue | orange | orange | blue | blue | orange | blue |
| trans P-A V2 | blue | blue | blue | blue | blue | blue | blue | orange | blue | blue | orange | orange | blue | blue | blue | orange | blue | blue | blue | blue | orange | blue |
| trans P-A V3 | orange | blue | blue | orange | orange | blue | orange | blue | blue | blue | orange | orange | blue | blue | blue | blue | blue | blue | orange | blue | orange | blue |
| trans P-A V4 | blue | blue | blue | blue | blue | orange | orange | blue | blue | blue | orange | blue | orange | blue | blue | blue | orange | blue | orange | orange | blue | blue |
| trans P-A V5 | blue | blue | blue | blue | blue | blue | blue | blue | blue | orange | orange | blue | blue | blue | blue | blue | blue | blue | blue | orange | blue | blue |
| trans A-P V1 | orange | blue | orange | blue | orange | blue | orange | blue | blue | orange | orange | blue | blue | orange | blue | blue | orange | blue | blue | orange | blue | blue |
| trans A-P V2 | blue | orange | orange | blue | blue | orange | blue | orange | blue | orange | blue | blue | orange | orange | blue | orange | orange | orange | blue | orange | blue | blue |
| trans A-P V3 | blue | blue | orange | orange | blue | blue | blue | orange | blue | blue | orange | blue | blue | orange | blue | blue | blue | blue | orange | blue | blue | blue |
| trans A-P V4 | orange | blue | orange | blue | blue | blue | blue | orange | orange | orange | blue | orange | blue | blue | blue | blue | blue | blue | blue | orange | blue | blue |
| trans A-P V5 | blue | blue | orange | blue | orange | orange | blue | orange | blue | orange | orange | blue | blue | blue | orange | blue | orange | blue | blue | orange | orange | orange |
| comp P-A V1 | blue | blue | orange | blue | blue | orange | orange | orange | orange | orange | orange | blue | orange | blue | blue | orange | orange | blue | blue | blue | blue | blue |
| comp P-A V2 | orange | blue | orange | blue | blue | orange | orange | blue | orange | orange | orange | orange | orange | blue | blue | blue | orange | blue | blue | orange | orange | blue |
| comp P-A V3 | blue | blue | orange | blue | blue | orange | orange | blue | blue | orange | orange | blue | orange | blue | orange | orange | orange | blue | blue | orange | blue | blue |
| comp P-A V4 | blue | orange | blue | blue | blue | orange | orange | orange | orange | orange | orange | blue | orange | blue | blue | blue | orange | orange | orange | orange | blue | blue |
| comp P-A V5 | blue | orange | orange | blue | blue | orange | orange | blue | orange | orange | orange | blue | orange | orange | orange | orange | orange | blue | blue | orange | orange | blue |
| comp A-P V1 | blue | orange | orange | blue | orange | orange | orange | orange | blue | orange | orange | orange | orange | blue | orange | blue | orange | blue | orange | orange | orange | blue |
| comp A-P V2 | blue | orange | orange | orange | orange | orange | blue | orange | orange | orange | orange | orange | blue | blue | orange | blue | orange | blue | orange | orange | blue | orange |
| comp A-P V3 | blue | orange | orange | blue | blue | orange | orange | orange | orange | orange | orange | orange | orange | orange | blue | blue | orange | blue | orange | orange | blue | blue |
| comp A-P V4 | orange | orange | orange | blue | orange | orange | orange | blue | orange | orange | orange | blue | orange | blue | blue | orange | orange | blue | blue | orange | blue | blue |
| comp A-P V5 | orange | orange | blue | orange | blue | orange | orange | blue | blue | orange | orange | blue | orange | blue | blue | orange | orange | blue | orange | orange | blue | blue |

**Supplementary Table.3** Summary of dynamic factor analysis (DFA) models. For each trajectory, the DFA model was performed iteratively, starting with 1 common trend. The Akaike information criterion (AIC) and AIC corrected for small sample size (AICc) of each model are shown above. Since AICc can prevent over-fitting, it was used to determine the best-fit approximating model. For looming stimuli, the best model contains 7 common trends. For translational stimuli, the best model contains 3 common trends. For compound stimuli, the best model contains 5 common trends

| | **Loom_V1** | | **Translational_AtP_V1** | | **Compound_AtP_V1** | |
|---|---|---|---|---|---|---|
| | **AIC** | **AICc** | **AIC** | **AICc** | **AIC** | **AICc** |
| **CT** | 26558.54 | 26573.25 | 28898.23 | 28922.89 | 30942.57 | 30955.52 |
| **2 CTs** | 25897.78 | 25957.68 | 29650.71 | 29669.32 | 30471.55 | 30524.14 |
| **3 CTs** | 25164.5 | 25301.97 | 28875.01 | 28917.0 | 30060.52 | 30181.1 |
| **4 CTs** | 24524.79 | 24775.75 | 28977.04 | 29052.38 | 29725.07 | 29943.86 |
| **5 CTs** | 23974.08 | 24376.16 | 29094.94 | 29213.59 | 29502.02 | 29851.23 |
| **6 CTs** | 23765.19 | 24359.46 | 29201.69 | 29373.94 | 29431.48 | 29945.54 |
| **7 CTs** | 23487.60 | 24318.57 | 29301.48 | 29537.94 | 29354.46 | 30070.24 |
| **8 CTs** | 23216.88 | 24332.92 | 29396.26 | 29707.80 | 29312.40 | 30269.45 |
